# Streamlined generation of CRISPR/Cas9-mediated single-cell knockout clones in murine cell lines

**DOI:** 10.1101/2023.10.20.563250

**Authors:** Hub Tobias, Cornean Alex, Round Kellen, Fleming Thomas, Freichel Marc, Medert Rebekka

## Abstract

Clonal cell lines harbouring loss-of-function mutations in genes of interest are crucial for studying the cellular functions of the encoded proteins. Recent advances in genome engineering have converged on the CRISPR/Cas9 technology to quickly and reliably generate frameshift mutations in the target genes across various cell lines and species. Although high on-target cleavage efficiencies can be obtained reproducibly, screening and identifying clones with loss-of-function alleles remains a major bottleneck. Here, we describe a single sgRNA strategy to generate CRISPR/Cas9-mediated frameshift mutations in target genes of mammalian cell lines that can be easily and cost-effectively identified. Given the proliferation of workhorse cell lines such as HEK293 and N2a cells and the resulting clonal expansion of the cell type, our protocol can facilitate the isolation of knockout clonal cell lines and their genetic validation within a period of down to 3-4 weeks.

## Introduction

The forward geneticist’s toolkit has seen substantial progress in the 21^st^ century, with CRISPR/Cas systems emerging as the dominant approach to genome editing in basic research and gene therapy (1, 2). Originally discovered as an immune defence mechanism in *Streptococcus pyogenes* bacteria (3–6), the CRISPR-associated Cas9 protein couples with a guide RNA (gRNA) to identify the three nucleotide NGG protospacer adjacent motif (PAM) (7–9). Once detected, sufficient complementarity among the gRNA and target site allows the Cas9 enzyme to effectively introduce DNA double-stranded breaks (DSB) to particular genomic regions (7–12).

Introduced DSBs can be repaired by either error-free, albeit rarer, homology-directed repair (HDR) or end-joining of the obtained strands. Within eukaryotic cells, DSBs are predominantly repaired by end-joining processes such as non-homologous end joining (NHEJ) and microhomology-mediated end joining (MMEJ). While the cellular repair pathways are generally very efficient in perfectly restoring the initial DNA state, end processing of the two strands after DSBs is somewhat error-prone, eventually leading to an altered nucleotide sequence after ligation. Changes in the nucleotide sequence are mostly in the form of insertions or deletions, commonly referred to as indels (8). Introducing these indels frequently causes a shift in the reading frame that ends with a premature termination codon (PTC), and has been described as efficiently yielding knockout (KO) cells (8, 13).

Despite the wide-ranging benefits of the CRISPR/Cas technology for genome engineering in cells, identifying the optimal method for delivering the CRISPR/Cas components remains challenging (14, 15). The decision on the choice for delivery typically depends on the precise application and cell line used. In short, the CRISPR components can be delivered either as plasmids (7, 16, 17), as *in vitro* transcribed (IVT) or chemically synthesized sgRNA and IVT Cas9 mRNA (18), and via the assembly of Cas9 protein and sgRNA into ribonucleoprotein (RNP) complexes *in vitro* (19).

In addition to the type of cargo delivered, the delivery method is also critical for efficient genome editing. Given the low cost and universal access, lipid-based nanoparticle (LNP) delivery, such as lipofection of plasmids, is the predominant method to deliver CRIPR/Cas9 components (14). For some cell lines, however, lipofection might not effectively deliver the Cas/gRNA cargo and therefore, more versatile, albeit costly and technically much more challenging alternatives, such as electroporation and viral delivery, are required (14, 15).

Although several guidelines and protocols exist to aid the generation of CRISPR-based KO cells, these tend to fail to accurately characterise the genetic makeup of the isolated and expanded KO cell lines (16, 20, 21). Validating efficient genome editing events from transfected cells requires initial identification of putative alleles, commonly identified with the aid of designated algorithms such as “Interference of CRISPR Edits” (ICE) (22) that use standard Sanger sequence traces to provide a profile of editing efficiency. However, these tools used in isolation cannot accurately determine the actual allele frequencies and are limited by their inability to detect longer indels in the target gene (23). Targeted deep sequencing has recently become more affordable for genome editing validation further allowing detailed and accurate sequence analysis to supplement initial, cost-effective Sanger sequencing-based screening.

Here, we present a fast and cost-effective approach for the generation and molecular characterization of knockout cell clones by CRISPR/Cas9-mediated gene deletion. From cell transfection and clonal expansion to final genetic frameshift verification, this study aims to describe a comprehensive workflow for generating mammalian knockout cell lines.

## Materials and methods

### Cell culture

#### N2a cells

The murine neuroblastoma cell line N2a (DSMZ, ACC 148) was cultured in DMEM-GlutaMAX^TM^ (Gibco^TM,^ Thermo Scientific) supplemented with 10% FCS (v/v) and 1% Pen/Strep (v/v) at 37°C in a saturated humidity atmosphere containing 5% CO_2_. Cells were grown to 70% confluence for transfection with CRISPR/Cas9-plasmid constructs and passaged every 3-4 days below 80% confluency using trypsin digest at 37°C for 3 min.

#### MCE cells (MCECs)

Murine cardiac endothelial cells (MCEC) were obtained from Biozol/CELLutions Biosystems Inc (Catalogue No. CLU510). Cells were cultured in DMEM, low glucose, pyruvate (Gibco^TM,^ Thermo Scientific, 31885-023) supplemented with 5% FCS (Sigma), 1% Pen-Strep (Sigma), 1% Amphotericin B (Sigma) and 1% HEPES (Sigma) in a humidity incubator at 37°C and 5% CO_2_. Prior to cell seeding culture flasks were coated with 0.5% gelatin (w/v, Sigma) in DPBS (Gibco^TM,^ Thermo Scientific) for 15 min at RT. MCEC were split every 2-3 days using TrypLE express enzyme (Gibco^TM,^ Thermo Scientific) for 5min at 37°C. Transfection was performed at 70% confluence.

#### sgRNA cloning

Specific sgRNA protospacers sequences (Table 1) were ordered from Sigma Aldrich as oligonucleotides (Table 2) and were cloned into pSpCas9(BB)-2A-GFP (PX458, a gift from Feng Zhang, Addgene plasmid #48138; http://n2t.net/addgene:48138; RRID: Addgene_48138) containing pSpCas9 and an eGFP reporter cassette. sgRNAs were cloned as described previously (17)). sgRNA plasmids were transformed into chemically competent *DH5α* and selected on LB plates with ampicillin (100 µg/ml) by incubation at 37°C for 16-18 hours. Selected bacterial clones were cultured in LB medium with ampicillin (100 µg/ml) and incubated overnight at 37°C and 220 rpm. Plasmids were extracted using the ZymoPURE^TM^ Plasmid Miniprep kit following the manufacturer’s instructions. Correct plasmid sequences were validated by Sanger sequencing (Eurofins Genomics) using the primer listed in Table 2. Transfection-grade plasmids of sequence-validated clones were obtained using the EndoFree Plasmid Maxi Kit (Qiagen) according to the manufacturer’s protocol.

**Table 1.**
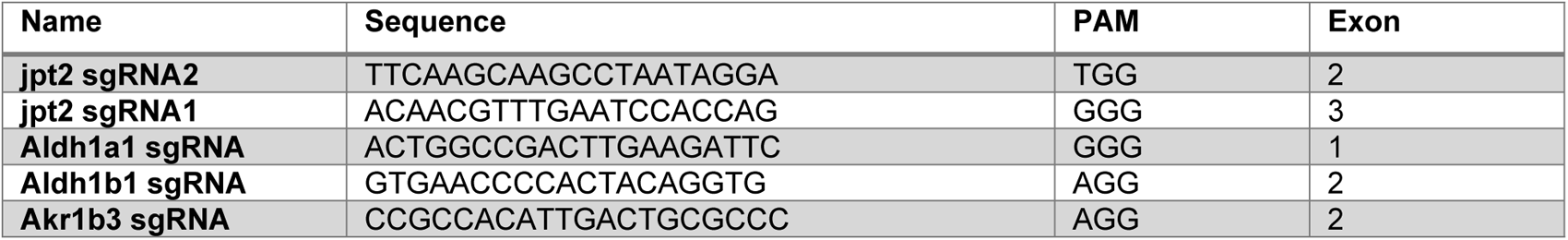
sgRNAs used in this study.

**Table 2.**
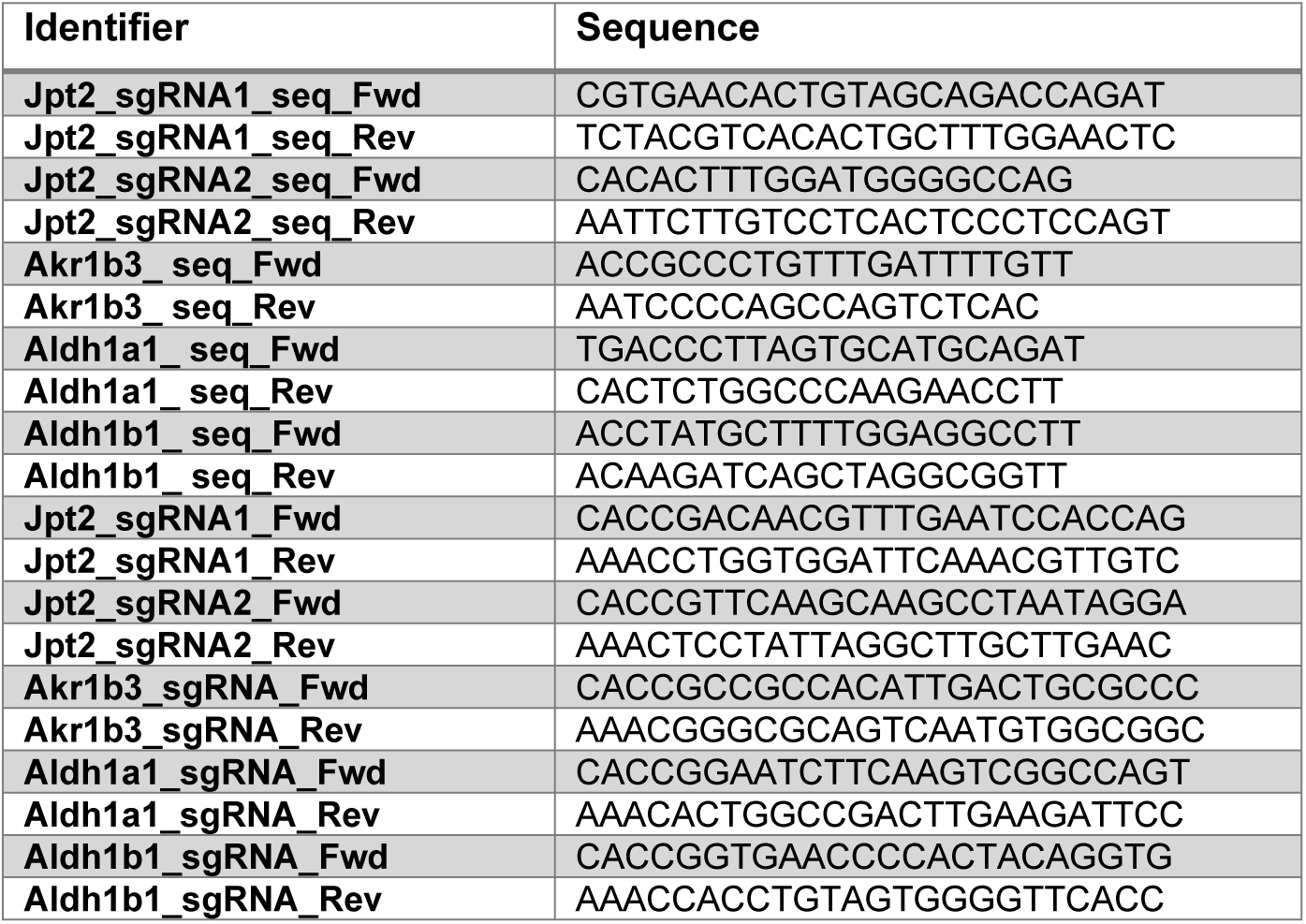
Oligos used for genotyping (_seq_) and sgRNA cloning (_sgRNA_).

### Transfection

#### N2a cells

Transfection of N2a cells was performed in a 10 cm petri dish using Lipofectamine RNAi MAX (Thermo Scientific) according to the manufacturer’s protocol. Therefore 1.3 x 10^6^ cells were plated 24 hours prior to transfection on a 10 cm culture dish. Transfection solution was prepared by mixing 43 µl RNAi MAX with 500 µl Opti-MEM^TM^ (Thermo Scientific) and separately 14.5 µg CRISPR/Cas9 plasmid containing the cloned protospacer with 500 µl Opti-MEM^TM^ (Thermo Scientific). The final transfection solution was obtained by mixing the prepared solutions. Cells were transfected by adding 1 ml of transfection solution to the plated cells, cultured in 10 ml DMEM+GlutaMAX^TM^ (Thermo Scientific) + 10% FCS. Cells were incubated in transfection solution for 48 hours prior to cell harvest and preparation for FACS sorting.

#### MCE cells (MCECs)

MCECs were transfected via electroporation using the Neon^TM^ transfection system (Invitrogen^TM^, Thermo Scientific). Therefore, cells were harvested, counted and 1.5x10^6^ cells resuspended in 150 µl of resuspension buffer R containing 15 µg CRISPR/Cas9 plasmid containing the cloned protospacer. Electroporation parameters were selected as followed: 1200V, 30 ms Pulse with, 1 pulse. After electroporation cells were immediately transferred to a 0.5% gelatine coated culture dish filled with 10 ml medium. For optimal growth, medium was changed 24h after transfection.

### FACS sorting

48 hours after transfection, 50% of cells were used for single-cell sorting. Therefore, cells were harvested and re-suspended in 2 ml DPBS (Thermo Scientific), supplemented with 10% FCS, 1% Pen-Strep and 1% Amphotericin B (Sigma) to obtain a single cell suspension. Cells were sorted for GFP reporter expression into two 96-well plates. The remaining GFP-positive cells were batch sorted, resulting in a GFP+ cell pool for further gene editing analysis. FACS sorting was performed using a FACS Aria III (BD). The remaining 50% of cells were used for DNA analysis (cell pool analysis).

### DNA preparation for gene editing analysis

#### DNA extraction

For cell pool analysis, 50% of transfected cells were transferred to a fresh 15 ml falcon spun down and re-suspended in 2 ml DPBS which was transferred to a 2 ml Eppendorf tube. Cells were again spun down and supernatant was discarded, following genomic DNA extraction using 200 µl Direct PCR (Ear) lysis reagent (Viagene), supplemented with 0.5 µg Proteinase K (AppliChem). The mix was incubated for 15 min at 55°C following 15 min at 85°C. To investigate editing efficiency in the single cell clone-derived colonies, genomic DNA was extracted using 50 µl of Direct PCR (Ear) lysis reagent (Viagene), supplemented with 125 ng Proteinase K (AppliChem) directly in the well of the 96 well plate. Cells were re-suspended in lysis buffer transferred to a 2 ml Eppendorf tube and incubated at 55°C for 15 min followed by 85°C for 15 min.

#### Sanger sequencing and analysis of PCR amplicons

Primer pairs used to amplify the locus around the target site are listed in Table 2. PCR was performed at 30 cycles using Q5 High fidelity DNA Polymerase (NEB) at 0.02 U/µl according to further manufacturer’s instructions. The specific PCR product was separated on a 1% agarose gel by gel electrophoresis, the band excised and purified using the QIAquick Gel Extraction Kit (Qiagen) following the manufacturer’s protocol. Purified PCR amplicons were Sanger sequenced (Eurofins Genomics) using the forward primer listed in Table 2. Resulting Sanger traces were analysed using the ICE tool (Synthego, https://ice.synthego.com/#/ (24)).

#### Targeted amplicon sequencing

N2a clones assessed by ICE analysis were further validated by targeted deep-sequencing. PCR was performed as described above and PCR products were purified using column-based purification (innuPREP PCRpure kit, Analytic Jena). The sample concentration was determined using the Qubit dsDNA BR Assay Kit on a Qubit 4 Fluorometer (Thermo Fisher Scientific). Purified PCR products adjusted to 25 ng/µl were sequenced using the Amplicon-EZ service at GeneWiz (Azenta Life Sciences) on an Illumina MiSeq (2 x 250 bp sequencing, paired-end).

### Analysis and data visualisation

#### Analysis and plotting of NGS data

Amplicon sequencing data were analysed with CRISPResso2 v.2.2.11 in CRISPRressoBatch mode in conjunction with gene_input.txt files that were populated with the following parameters: the respective --fastq_r1 (r1) and --fastq_r2 (r2) files present in the given folder, the target amplicon sequence (a), sgRNA sequence (g) and the names of the output file (N), folder (o) and the min_average_read_quality (q) set to 30 (25). The following files generated file generated by CRISPResso2 were then used to generate the depicted HTS figures / tables: “Nucleotide_percentage_quilt_around_sgRNA_(…).pdf” (summarising all samples per gene), “Alleles_frequency_table_around_sgRNA_(…).pdf” (for the visualisation of allele composition in each clone), and “Alleles_frequency_table_around_sgRNA_(…).txt” for further calculations (regarding each clone), which were conducted as follows: 1) To remove PCR/Sequencing artefacts, the column %Reads was filtered as “Greater or Equal to 1.5” in each of the .txt files in MS Excel. 2) The filtered values were then copied to a new table and used to normalise after filtering, by adjusting to the total number of reads that is present for each filtered clone, as such #Reads[Allele1]/#Reads[Sum(Allele1…n, if >1.5perc)]*100.

#### Flow cytometry analysis

Flow cytometry data were analysed with FlowJo software v.10 (BD Biosciences).

#### Analysis and data visualisation

Data from different assays were collected in MS Excel. Graphs and statistical analyses were created using Prism 9 (GraphPad). Some icons and schemes were created using www.biorender.com (26). Figures were assembled in Adobe Illustrator.

## Results

### A fast and efficient workflow to establish mutant cell lines

Here we propose a six-step protocol to generate and identify single-cell clones with bi- or multiallelic frameshift mutations (**Fig 1**). From design to validation, this execution of the workflow will take between 8-12 weeks yielding stable and NGS-validated cell lines.

**Fig. 1.**
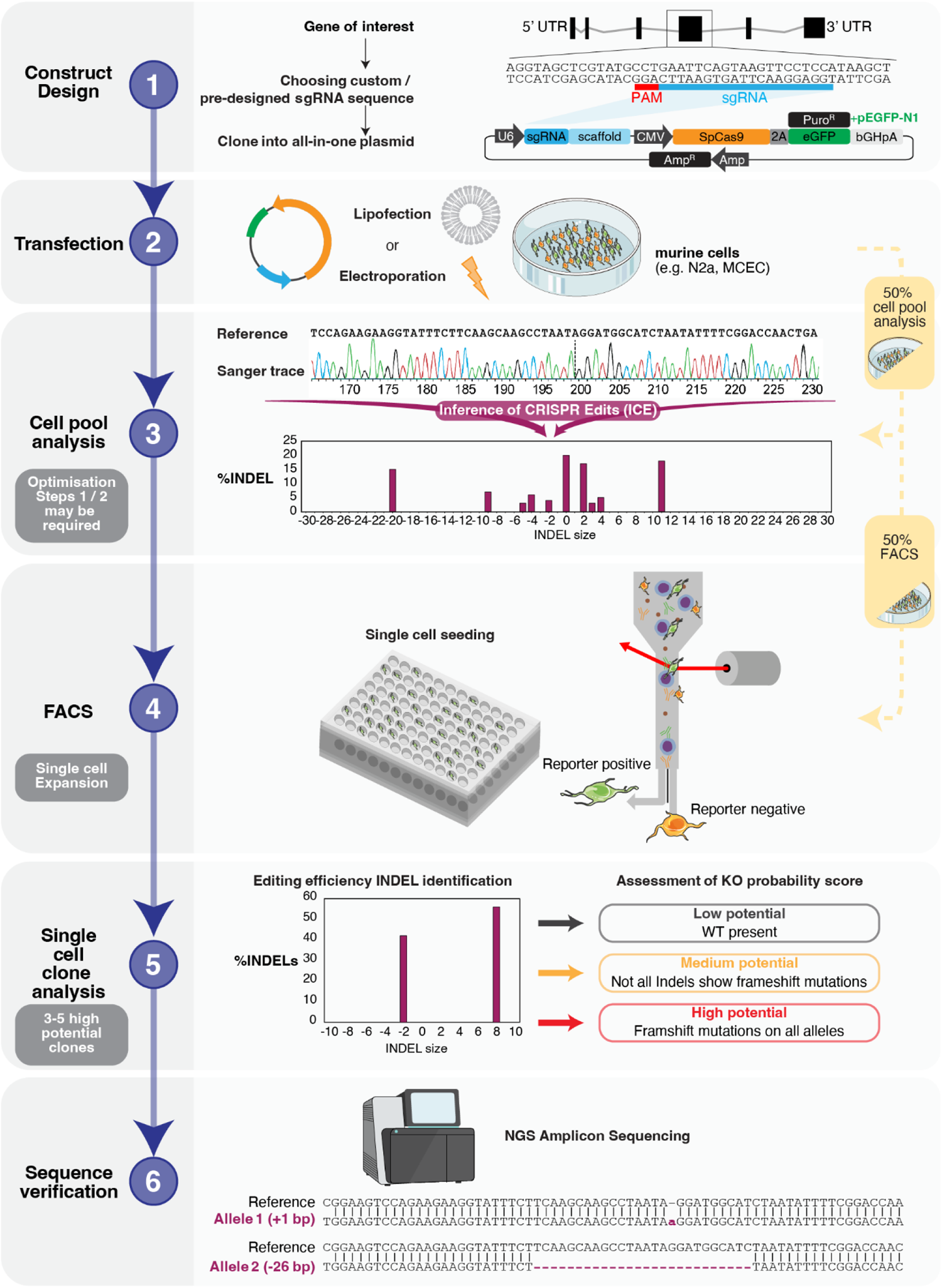
Stepwise workflow for generating single-cell knock-out (KO) clones with frameshift mutations. **Step 1**: Design of CRISPR construct. Illustration of target locus with location and sequence of target-specific gRNA (blue). An all-in-one plasmid with all components serves as the backbone for cloning sgRNA. **Step 2**: Transfection of the target cell line. **Step 3**: A fraction of the transfected cells is used for cell pool analysis. PCR products covering the editing site were generated, and Sanger sequenced, followed by Inference of CRISPR Edits (ICE) analysis. **Step 4**: The other fraction of transfected cells was used for FACS-based single-cell sorting of GFP+ cells. **Step 5**: Single-cell clone analysis and categorisation into low, medium and high potential clones based on editing and knockout scores. **Step 6**: Sequence validation of up to five high-potential clones using NGS sequencing allows precise sequence verification around the Cas9 cut site.

#### Step 1: Design and cloning of CRISPR construct

CRISPR sgRNA design and prediction tools that help to select target-specific sgRNAs have become ubiquitous. These can be either pre-designed (e.g. IDT, (27)) or customised sgRNAs via online prediction tools (e.g. CCTop, (28); CRISPRscan, (29); CHOPCHOP, (30); Synthego, (24)). Once a target-specific sgRNA has been selected, the primers to analyse target-derived alleles must be designed. In addition to standard primer design practices, we recommend to space primers for Sanger sequencing at least 100 bp 5’ of the sgRNA target site and at least one of the primers at a maximum distance of 600 bp. To validate final clones by amplicon deep-sequencing, it is also best to keep the full amplicon size <400 bp.

To clone the sgRNAs into the sgRNA-Cas9 all-in-one plasmid (such as pSpCas9(BB)-2A-GFP) expressing both Cas9 and a fluorescent reporter gene (e.g. GFP) separated by a 2A peptide and the U6 promoter for sgRNA expression, order sgRNA specific oligos as previously described (17). In brief, order top and bottom complementary oligos that contain the protospacer sequence with 5’ BbsI overhangs, and keep in mind to add a Guanosine at the 5’ end of the protospacer sequence (to ensure efficient transcription form the U6 promoter), in case the target site does not start with a G (PAM-distal end). Following oligo annealing, ligation and transformation of *E. coli,* final plasmid sequences are confirmed by Sanger sequencing.

#### Step 2: CRISPR/Cas9 construct delivery

Delivering the designed and cloned plasmids is most easily accomplished by standard lipofection, using Lipofectamine reagents (RNAi MAX, 2000, 3000 etc.), which we used to transfect the workhorse N2a cell line. There are, however, difficult-to-transfect cell lines, in our case, the MCECs, for which electroporation using any of the available commercial systems (e.g. 4D Nucleofector [Lonza], Neon transfection system [Thermo]), is recommended. Importantly, our workflow does not require any optimisation and has been used to establish mutant lines with transfection efficiencies as low as 12%.

Following plasmid delivery, cells are cultured for 48 hours to allow for strong expression of specific reporter proteins. The entire culture dish is then harvested, and cells are resuspended to obtain a single-cell suspension. Of these cells, half are directly used for editing (cell pool) analysis. The other half of the cell pool is used for immediate FACS single-cell sorting.

#### Step 3: Cell pool analysis and editing control

To assess the extent of gene editing in the transfected cell pool, the target locus is amplified and sequenced with standard Sanger chemistry. The resulting Sanger sequencing traces are then analysed using the ICE online tool (Synthego) (22), which can deconvolute mosaic traces and provide information about indel occurrence, length and frequency and thereby estimate overall editing efficiency. Here, editing efficiency refers to the percentage of non-wild-type sequences in the pool.

If genome editing is not observed in the cell pool, the sgRNA design and transfection parameters in Step 1 and Step 2 must be optimised. Typically, it is sufficient to design and clone an alternative sgRNA to overcome editing difficulties.

#### Step 4: FACS single-cell sorting of KO clones

The GFP reporter expression of the all-in-one plasmid can be used for fluorescent-based single-cell sorting to enrich cells that express the Cas9/sgRNA-expressing plasmid. Reporter-positive single cells are sorted onto one to two 96-well plates. In cases where you may have obtained alternative sgRNA plasmid constructs without an EGFP reporter cassette, we recommend supplementing the delivery mixture of plasmid with a small quantity of an EGFP-containing plasmid to facilitate the enrichment of transfected cells. In these instances, however, some EGFP-positive cells may not express the Cas9/sgRNA components.

#### Step 5: Expansion and single-cell clone screening

Single cells are then cultured and expanded for 1-2 weeks, depending on the proliferation rate of the cell line. To identify and assess edited clones, a fraction of cells is used to isolate genomic DNA at the earliest possible time at which cells can be passaged. Next, target regions are PCR amplified and analysed by Sanger sequencing and ICE deconvolution. Here, we introduce three categories by taking the presence of WT alleles, the indel size and the potential zygosity into account. The latter considers the number of edited alleles in relation to the total number of alleles identified during ICE analysis. Cell clones which still display a WT sequence after editing are considered as low potential. The ICE algorithm defines clones without WT sequence either as having intermediate potential if not all indels meet the requirements for a knock-out mutation (not multiple of three or <21 bp) or as high potential if all identified indels, lead to a frameshift (not multiple of 3) or exceed an indel size of 21 bp (24). Here, we follow the pre-set parameters of the ICE algorithm. To ensure the selection of promising clones, an individual assessment of the size of the deletion (>21 bp) is recommended.

#### Step 6: Sequence verification

Once 3-5 high-potential clones are identified using ICE analysis, sequence validation can be performed using Next Generation Sequencing (NGS) of target locus PCR products. To this end, PCR products are sequenced on an Illumina-based sequencing platform (e.g. MiSeq) which yields detailed insights around the Cas9 cut site, thus enabling verification of cell clones with frameshift mutations on all alleles of the target gene.

#### Troubleshooting

We have compiled a table of issues that we and others have encountered, accompanied by the most likely underlying reason and recommended solutions in Table 3.

**Table 3.** Troubleshooting of common problems faced during workflow establishment.

| Problem faced | Possible reason | Solution |
| --- | --- | --- |
| Poor Sanger sequencing quality | DNA impurity and / or low/high concentration<br>Large deletion close to the site of primer annealing | Optimize DNA extraction method and submit appropriate concentration for sequencing<br>Use an agarose gel extraction protocol (instead of PCR clean-up)<br>Try sequencing from other direction<br>Design primer with a minimum of 150 bp up- and downstream of Cas9 cut site |
| Poor quality of ICE results | Poor quality of Sanger traces<br>Large deletions | Increase Sanger quality as described above<br>Have a look at the ICE-D values provided in the download files as ICE only addresses INDEL < Δ 30 bp<br>Always process a WT sample along to ensure right genetic background for alignment |
| **ICE warning: “guide not found in control sequence”** | • Incorrect spelling or orientation of sgRNA during<br>• Single nucleotide polymorphism (SNP) within the cell line compared to the reference genome | • Ensure 5’-3’ orientation<br>• Ensure that you are on the right strand<br>• Check the Sanger trace for mutations compared to the designed sgRNA<br>• Always process a WT sample to ensure the right genetic background for alignment |
| **PCR reaction generates two bands in single-cell clones** | • Large INDELs<br>• Impurities<br>• Issues at the stage of single cell sorting | • Repeat PCR<br>• Run gel for extra time and analyse both bands independently<br>• Be aware that within a cell line there is a tendency towards similar INDEL sizes!<br>• Re-sort the clone and expand |

### CRISPR/Cas9 strategy to generate frameshift mutations in the *Jpt2* gene

To exemplify the proposed workflow, we targeted the *Jpt2* (synonym *Hnl1*) gene locus based on the mouse genome reference GRCm39 (GCA_000001635.9) (**Fig 2A**, **S1A Fig**). We targeted exon 3 (sgRNA#1) and exon 2 (sgRNA#2) of the *Jpt2* locus and designed primer pairs as described above. The gRNAs were cloned into the pSpCas9(BB)-2A-GFP (PX458) plasmid backbone (**Fig 2A**), resulting in two target site-specific all-in-one plasmid constructs. Sequence integrity after cloning was validated by Sanger sequencing of purified plasmid preparations. Following the transfection of individual all-in-one plasmids, their successful delivery of the construct was confirmed by fluorescence imaging (**Fig 2B**) without further consideration of transfection efficiency. Interestingly ICE cell pool analysis of unsorted N2a cells found no evidence for indels with gRNA#1, whereas gRNA#2-based Cas9 cleavage resulted in an indel score of 7% and a knockout score of 6% (**S1B Fig**). Importantly, our alignment of sequences obtained from the locus PCR product of exon 3 with the reference genome (mouse GRCm39) revealed that our N2a cells exhibit a single nucleotide polymorphism (SNP) in our designed gRNA#1 binding site region of the *Jpt2* gene, that is not present in the reference sequence we used for the gRNA design (**S1C Fig**), which stresses the point of sequencing target loci for every single cell line.

**Fig. 2.**
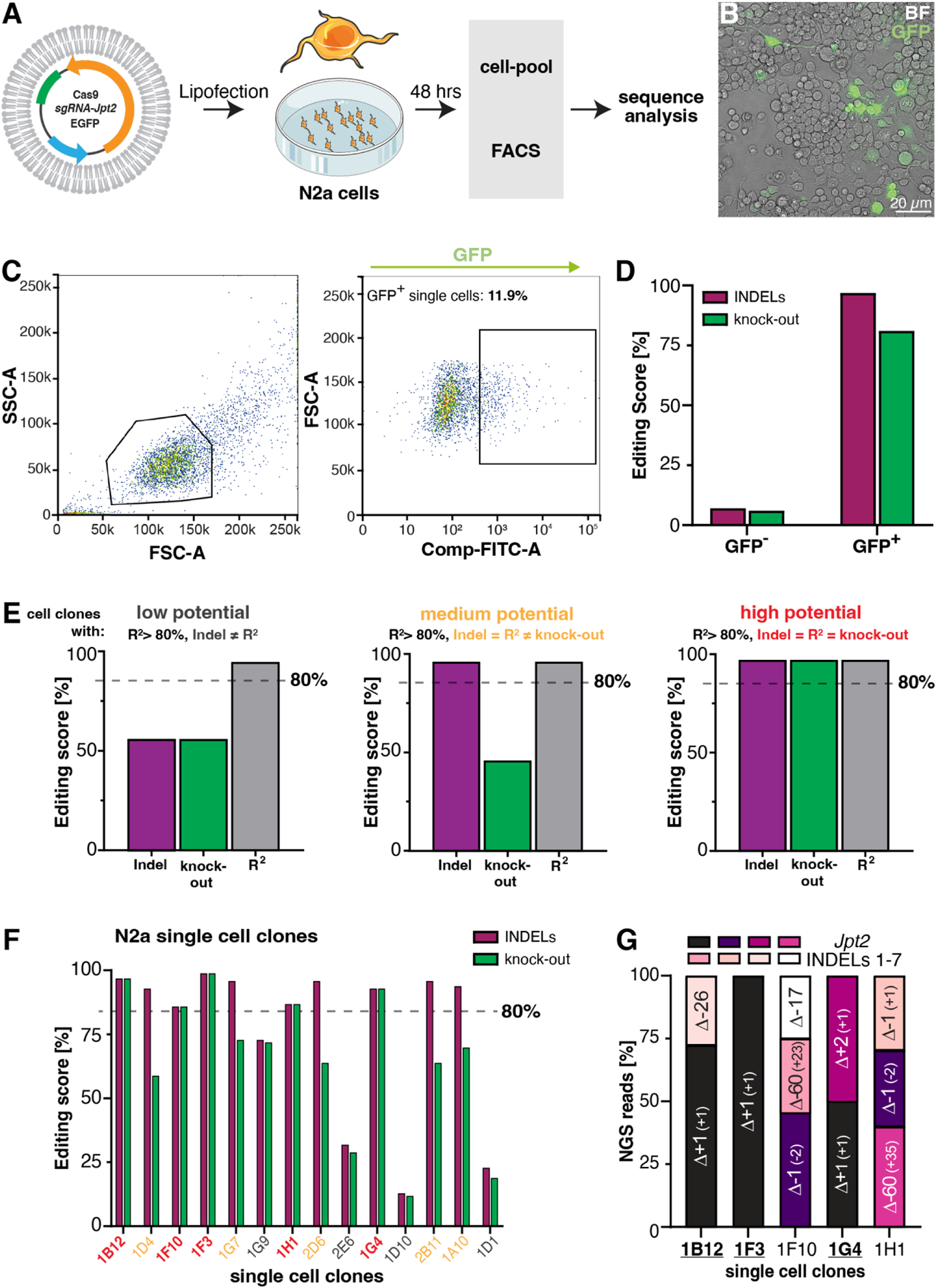
Straightforward generation of KO clones in murine N2a cells at the *Jpt2* locus. **(A)** Experimental setup for generating a *Jpt2* KO in N2a cells using an all-in-one plasmid PX458 carrying Cas9, sgRNA and GFP marker. Plasmid delivery is achieved using lipofection, followed by FACS single-cell sorting 48 h after transfection and sequence analysis. **(B)** Fluorescence image of N2a cells 48 h after lipofection with sgRNA#2 confirmed transfection success. **(C)** FACS analysis and sorting of single GFP+ cells for analysis and clonal expansion. **(D)** Indel and knock-out score obtained by ICE-based Sanger trace analysis of the GFP+ sorted cell pool (R^2^ > 0,95). **(E)** Categorization of single cell clones based on indel spectrum analysis. The R^2^-value serves as a quality score for the indel distribution proposed by ICE. Clones with a low-quality score (>80%) were not used for analysis. The knock-out score refers to indels resulting in a frameshift (Δ ≠ n*3 for n = ℤ) or indels larger than 21 bp. Here, clones with proposed WT sequences within the Sanger trace are considered low potential. Clones with a high indel score but reduced knock-out score are considered medium potential. Clones are classified as high potential if no WT sequences are identified and the knock-out score meets the indel score, which means that all proposed insertions or deletions are frameshift mutations. **(F)** ICE analysis of 14 single-cell clones revealed five high-potential clones (1B12, 1F10, 1F3, 1H1, 1G4). **(G)** Illumina amplicon-sequencing of all five high potential clones confirmed ICE-based categorisation as all single cell clones display at least 98% of Jpt2 knock-out mutant reads.

Consequently, all following experiments were performed with sgRNA#2. FACS sorting of GFP^+^ N2a cells after transfection revealed a transfection efficiency of 12% and allowed single-cell sorting into two 96-well plates to generate and expand isogenic single-cell clones (**Fig 2C**). We here compared GFP^+^ and GFP^-^-cell pool ICE analysis, showing a more than 13-fold increase (92%) of editing efficiency in the GFP+ pool (**Fig 2D, S2 Fig**). These results show that even given a low transfection efficiency and a respective low cell pool editing efficiency of 7%, FACS sorting can enrich edited cells, yielding an editing efficiency of 92% and a knockout score of 77% in the sorted pool. After clonal expansion for 2 weeks, 14 colonies were selected and analysed for frameshift mutations using ICE analysis. To estimate the knock-out potential of our clones, we here introduce three categories (**Fig 2E**), based on the ICE R^2^ score (R^2^) threshold of 80%, to preselect low-quality clones before detailed analysis. It is worth mentioning that R^2^ limits the maximum values indel and knockout scores can reach. Hence a clone in which R^2^ equals the indel score is considered fully edited. Clones were assessed as low potential, intermediate potential or high potential concerning the probability of carrying frameshift mutations at the target locus.

Screened clones were categorised based on the following characteristics: Low potential – clones with indel score that did not reach the maximum R^2^ value (indel score ≠ R^2^) (**Fig 2E** left panel). Thus, the indel distribution provided by ICE proposes the appearance of the WT sequence in the Sanger trace. Medium potential – clones which showed no signs of a WT sequence (indel score = R^2^) but showed a smaller knockout score than indel score (indel score > knock out score) (**Fig 2E** mid panel). Hence, medium potential clones carry at least one allele with an indel mutation that keeps the sequence in frame. Clones with frameshift mutations (indel: Δ ≠ n*3 or exceed the size of 21 bp) on all alleles are considered to have a high potential. These clones display the same editing score value for indel and knock-out and are most promising for downstream analysis. All 14 N2a single-cell clones obtained after clonal expansion were screened and categorised according to the previously introduced specifications (**Fig 2F**). Interestingly, following ICE analysis, we identified only one clone (1D1) carrying a WT sequence, while the remaining clones displayed only edited traces (**S3 Fig**). Clones 1G9, 2E6, 1D10 and 1D1 did not pass our quality control of R^2^>80% and were excluded from the NGS analysis. We identified the clones 1B12, 1F10, 1F3, 1H1 and 1G4 as high potential knock-out clones according to their indel and knock-out scores (in red, **Fig 2F**).

We next analysed the high-potential clones by amplicon next-generation sequencing. The obtained sequencing data (**Fig 2G**, **S4 Fig**) revealed that all five clones carry only frameshift indel mutations at the target site and were correctly categorised as high-potential knockout clones. Surprisingly, indels were not distributed in a 1:1 ratio as expected for a diploid genome. For clone 1B12, we found a Δ-26 allele in 26% of the sequence reads (115,393), and a Δ+1 allele in 72% of reads. For clone 1F3, only one type of deletion allele, Δ+1, was identified in 95% of the reads (118,416). For clone 1F10, three indels alleles, Δ-1 (46%), Δ-60 (29%) and Δ-17 (25%) were identified. We further identified indel alleles Δ+1 (50%) and Δ+2 (49%) in clone 1G4 as well as indel alleles Δ-60 (39%) and Δ-1 (61%) in clone 1H1. Interestingly we identified a large deletion of Δ-60 bp in NGS data of clones 1F10 and 1H1, which was not identified by ICE analysis (**S3 Fig**). However, it is worth noting that ICE analysis (**S3 Fig**) and NGS sequencing (**S4 Fig**) showed similar indel distributions for the clones. Taken together, we demonstrate that the generation, isolation and validation of CRISPR/Cas9-based knockout clones is feasible in mammalian cell culture with minimal optimisation effort while maintaining a high standard of precision. Pairing FACS-mediated enrichment of transfected cells with a Sanger sequencing/ICE analysis-based pre-screen of clones followed by a final validation step using NGS sequencing combines speed with precision.

### Independent Generation of *akr1b3*, *aldh1a1* and *aldh1b1* knockout cell lines in mouse cardiac endothelial cells

We targeted three independent genes in mouse cardiac endothelial cells (MCECs) to test the established analysis workflow in a separate cell line. Here, we used a slightly different transfection strategy, using an all-in-one CRISPR plasmid without a fluorescence marker. We co-delivered with an EGFP plasmid using the Neon^TM^ transfection system. The downstream analysis steps remained unchanged (**Fig 3A**). With the two-plasmid approach and electroporation, we obtained considerably higher transfection efficiencies compared to transfection in N2a cells (**S5 Fig**). However, it is important to consider that with this approach GFP signal does not necessarily indicate the presence of the CRISPR plasmid in these GFP^+^ cells. ICE-based screening of *akr1b3* targeted single cell-based colonies revealed only one clone (3A2) meeting the previously set quality scores of R^2^=80% (**Fig 3B**). Due to a lack of high-potential clones, we reduced the quality score in this specific case to 75%, yielding four high-potential clones (3A2, 3B6, 3D3, 3B9). For *aldh1a1* and *aldh1b1,* we used a threshold of 80% and obtained four high-potential clones, respectively (**S6 Fig**). Of all available high-potential clones, we proceeded with three clones per target gene to validate by NGS sequencing (**Fig 3C**).

**Fig. 3.**
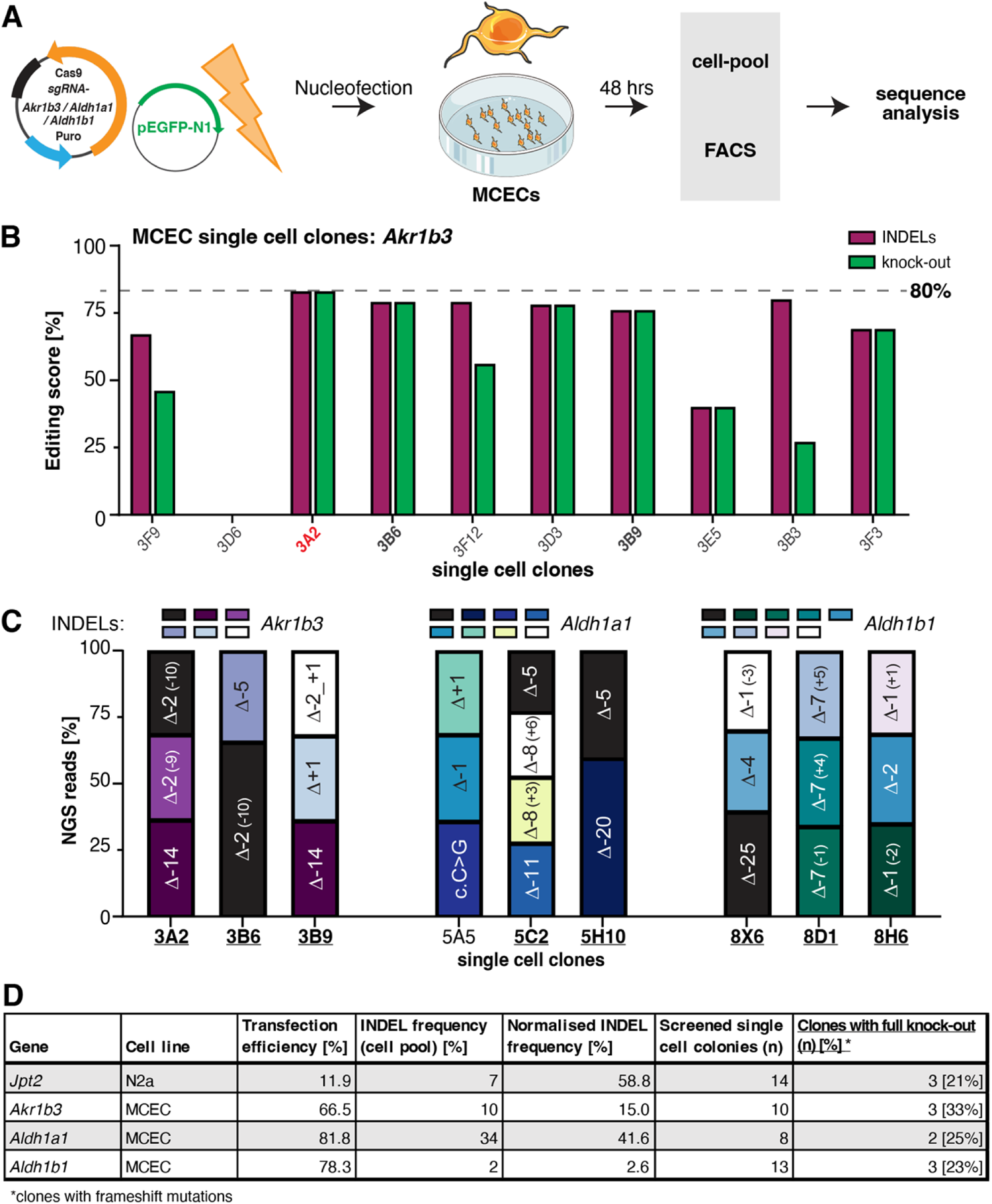
Application of the established workflow to targeting three independent genes in murine cardiac endothelial cells. **(A)** Experimental setup to generate *Akr1b3*, *Aldh1a1* and *Aldh1b1* KO MCEC cell lines using two plasmids, one carrying the Cas9 construct the other serving as GFP marker for single cell sorting. Plasmid delivery is achieved using nucleofection, followed by FACS single-cell sorting 48 h after transfection and sequence analysis. **(B)** ICE analysis of 10 single-cell colonies after targeting the *Akr1b3* locus in MCECs. Only one clone met the required previously introduced quality score of 80% (red). Due to a lack of clones meeting the quality score, we adjusted the quality score threshold for Akr1b3 to 75% resulting in 4 high potential clones, of which we used tree (3A2, 3B6, 3B9; bold) for further sequence analysis. **(C)** Illumina amplicon-sequencing results of all three targeted loci in MCECs. For all loci, three independent clones were analysed. **(D)** Summary of all independent genes targeted with the previously described workflow in MCEC and N2a cells. Normalised INDEL frequency was calculated by dividing the indel frequency by the transfection efficiency.

Surprisingly, clone 5A5 (*aldh1a1*) contained a single base change in one of the alleles and was therefore excluded. Our analysis of the remaining clones across all three target loci was similar to our ICE preselection analysis (**S8-S10 Fig**). Interestingly, for most clones, we observed three or four (clone 5C2) different alleles. Taken together, these results showed that ICE analysis of Sanger sequencing data allows precise preselection of high potential knockout clones that allows to reduce NGS sequence validation to a selected number of clones, reducing time and cost while obtaining substantial numbers of knockout clones (**Fig 3D**).

## Discussion

Our 6-step workflow describes a fast, reproducible and cost-effective means of creating and identifying knockout cell lines using a thorough understanding of the CRISPR/Cas9 technology (17, 31). For most cell lines, the optimal mode of delivery and the nature of the cargo itself will vary slightly (14, 15). Consequently, to identify ideal procedures and enrich properly edited cell lines that would facilitate clone identification, several rounds of optimisation are necessary, which unnecessarily prolong the mutant cell line generation process. Importantly, the combination of using FACS to enrich transfected cells with a robust pre-screening strategy of gene editing events in treated cells using Sanger sequencing and ICE analysis minimises the necessity to optimise transfection conditions. Given an all-in-one plasmid, that merely requires the design, selection and annealed oligo cloning of selected guide RNAs into Cas9-containing plasmid backbone, and which unlike mRNA or RNP approaches, can be used to enrich transfected cells, we have obtained knockout clones within a mere 4-6 weeks for four target genes.

The sorting of single GFP-positive cells is a crucial step for successfully expanding clonal knockout cell lines, preferentially used over transfected cell pools for downstream experiments, due to difficulties in readout reproducibility (32). The enrichment of successfully transfected cells is a key benefit of FACS sorting of GFP-positive cells following Cas9-linker-GFP transfections. In the case of our *Jpt2* knockout in N2a cells, this allowed to increase the editing efficiency from 7% in the unsorted pool to 92% in the sorted pool, thereby increasing the likelihood of obtaining knockout clones by a factor of 13. This decoupling from high transfection rates makes the workflow very versatile, with the need for costly transfection methods and labour-intensive optimisation steps of the transfection procedure.

The pre-screening step is essential for the increased speed in identifying suitable KO clones, and in our case, heavily relies on the ICE software to analyse Sanger sequencing reads. While other tools, such as the “Tracking of indels by Decomposition” (TIDE) tool (23) are similarly accurate in providing an estimate for the frequency of indels in the analysed sample, the ICE software is somewhat more accurate in correctly predicting the actual alleles. We have seen and validated this observation in all four genes targeted, where for smaller indels, the ICE analysis closely resembled our deep-sequencing analysis of amplicons. The indel analysis thereby reveals whether the clone still contains wild-type alleles (low potential), in-frame mutations (medium potential) or KO mutations on all alleles (high potential). However, NGS sequencing is necessary to confirm the KO mutations, identify larger indels omitted by the Sanger sequencing-based analysis and to determine the precise sequence of insertions at the target locus (33).

Isolating cell lines with biallelic frameshift mutations and the subsequent clonal expansion of single-cell progeny remain a significant challenge. The threat of off-target editing effects further corroborates this challenge. The generation of cell lines with any nuclease-based approaches, however, comes at a risk. DNA off-target editing remains a common problem (34), and while whole-genome sequencing has become more affordable, the analysis is a computational challenge for non-specialist labs. Alternatively, there are tools that predict off-target events, rank these and provide genome coordinates from where one can design primers to sequence these off-target sites (28, 35). And yet, in the case of identified off-targets, these cannot be removed from the cell clone’s genome by crossing as is the case when using mouse lines. Given that the probability of identical off-target events is low when examining multiple cell clones in which independent genome editing events achieved editing, the most straightforward solution to the off-target problem is to perform replicate experiments using several independent mutant alleles, ideally generated with different gRNAs.

It has been a frequently observed problem associated with immortalized cell lines, that these can differ in their polyploidy (36, 37). Therefore, if the cell line is not diploid more than two different alleles may be observed in these cells following CRISPR/Cas9-based genome engineering. Our finding of more than two alleles in N2a *Jpt2* KO cell clones and the indel distribution suggests that our N2a cells are polyploid. In accordance with this, N2a cells have previously been associated with polyploidy and a mean chromosome number of 102 (38). We additionally observed more than two alleles in most MCEC cell clones as well. Immortalization of this cell line has been achieved by lentiviral transfection of SV40 T antigen and human telomerase (Biozol/CELLutions Biosystems Inc). Importantly, SV40 T antigen immortalization is associated with chromosome instability and changes in karyotype (36, 37), which would explain the issue of allele number.

DNA sequence variants are very common in mammalian cells, including single nucleotide variants or SNVs. When using cell lines, these variants may pose a problem if the genetic makeup of these cells differs somewhat from the reference genome used. In our case, an SNV in the *Jpt2* locus of N2a cells resulted in a mismatch for one of the two gRNAs, completely abolishing nuclease-cleavage outcomes. Consequently, we recommend sequencing the target locus of your cell line of choice before designing your CRISPR/Cas9 experiment (Table 3).

A recent study investigated the correlation between Cas9-induced edits and protein loss, suggesting that cells with Cas9-induced frameshift most likely carry loss-of-function alleles (13), facilitating the creation of knockout cells. Although its cellular and physiological consequences have been only recently addressed, we now also know that transcripts containing PTCs may be subject to degradation by nonsense-mediated decay (39, 40). Consequently, frameshift mutations either lead to mRNA degradation with or without genetic compensation or give rise to severely truncated proteins (41). This is important in case the KO cell lines used for experimentation do not display any predictable loss-of-function behaviour and stresses the case for using several different clones carrying alternative allele combinations in downstream experiments.

In conclusion, our streamlined CRISPR/Cas9-KO approach enables the fast and cost-effective identification of KO cell clones with frameshift mutations. Depending on the cell proliferation and the resulting clonal expansion of the cell clones, KO cell lines can be generated and genetically validated within a period of down to 3-4 weeks.

## Acknowledgments

We thank Christin Richter, Hans-Peter Gensheimer, Beate Hilbert and Monika Langlotz from the Heidelberg University FACS core facility for expert technical assistance. We thank Feng Zhang for the plasmid pSpCas9(BB)-2A-GFP (PX458) (Addgene plasmid #48138). This research was supported by the Collaborative Research Centers (SFB) SFB1118 (project S03, projects A04 and S01 to TF), SFB 1328 (A21), SFB 1550 (project B09) of the Deutsche Forschungsgemeinschaft (DFG, German Research Foundation) and the DZHK (German Centre for Cardiovascular Research), the BMBF (German Ministry of Education and Research) (MF). RM is an alumnus of HBIGS, the Heidelberg Biosciences International Graduate School.

## Author contributions

<u>Tobias Hub</u>, Data curation, Investigation, Methodology, Visualisation, Writing – Original Draft Preparation, Writing – Review & Editing

<u>Rebekka Medert</u>, Conceptualization, Supervision, Formal analysis, Validation, Investigation, Writing – Original Draft Preparation

<u>Alex Cornean</u>, Data curation, Formal analysis, Visualisation, Writing – Original Draft Preparation, Writing – Review & Editing

<u>Kellen Round</u>, Methodology, Investigation, Writing – Original Draft Preparation

<u>Thomas Fleming</u>, Conceptualization, Funding acquisition, Resources, Writing – Review & Editing

<u>Marc Freichel</u>, Conceptualization, Funding acquisition, Project Administration, Resources, Writing – Review & Editing

## Data and materials availability

All data that support the findings of this study are available in this study within the manuscript and/or its supplementary materials.

## Supporting information

**S1 Fig.**
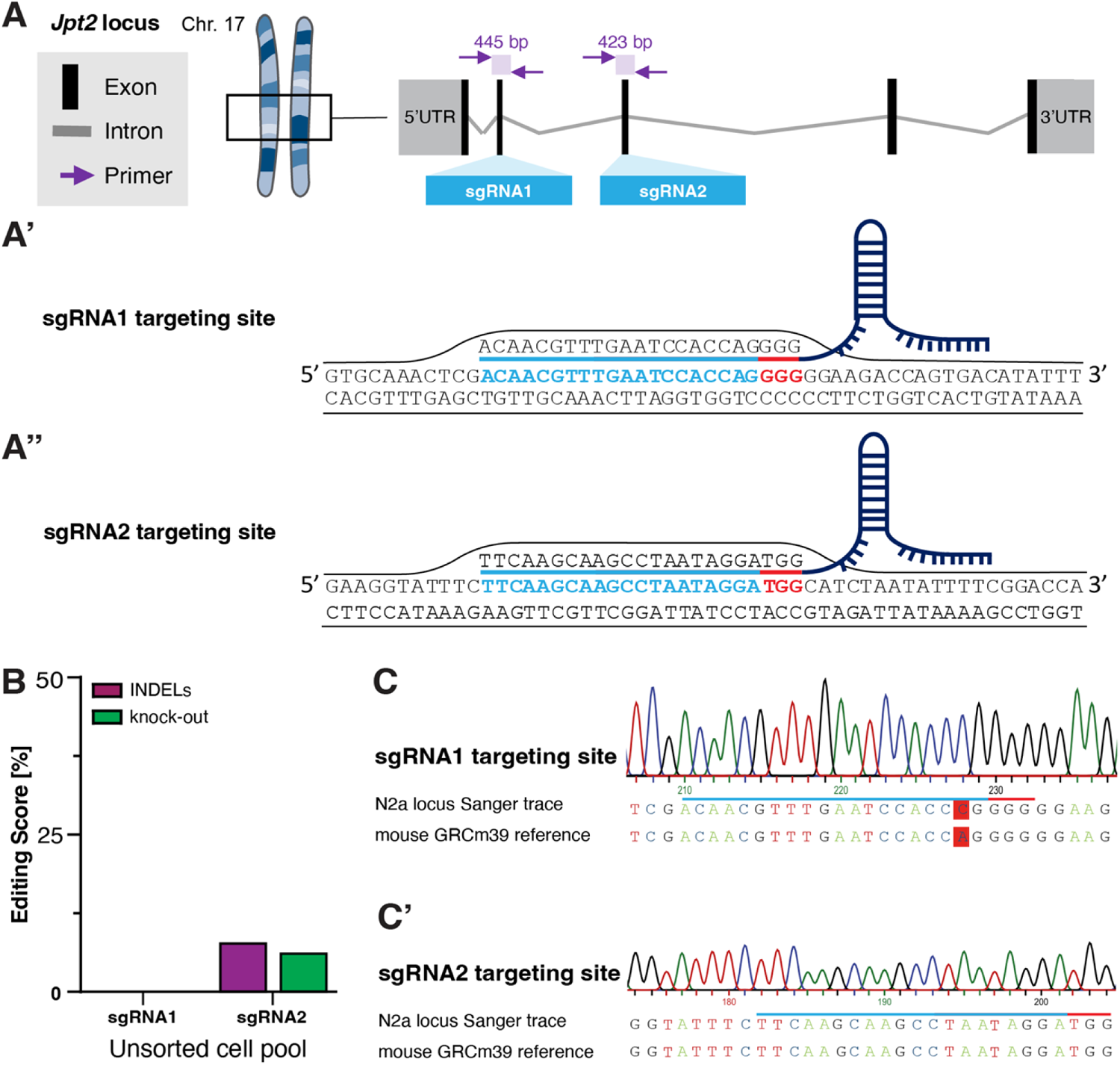
Targeting strategy for induction of frameshift mutations within the Jpt2 locus of murine N2a cells. **(A)** Graphical depiction of the genomic locus for *Jpt2* on mouse chromosome 17 with sgRNA#1 in exon 3 and sgRNA#2 in exon 2. (A’-A’’) Locus sequence information for sgRNA#1 (A’) and sgRNA#2 (A’’) target sites. The target-specific protospacer is highlighted in light blue and the NGG PAM motif is labelled in red. **(B)** Editing efficiency of both sgRNAs after ICE-based cell pool analysis (R^2^ > 0,95). **(C-C’)** Sanger traces reveal biallelic SNP (C -> A) in the used N2a genome compared to the reference sequence (ENSMUST00000024981_9 (Ensemble)), located in the sgRNA#1 binding site.

**S2 Fig.**
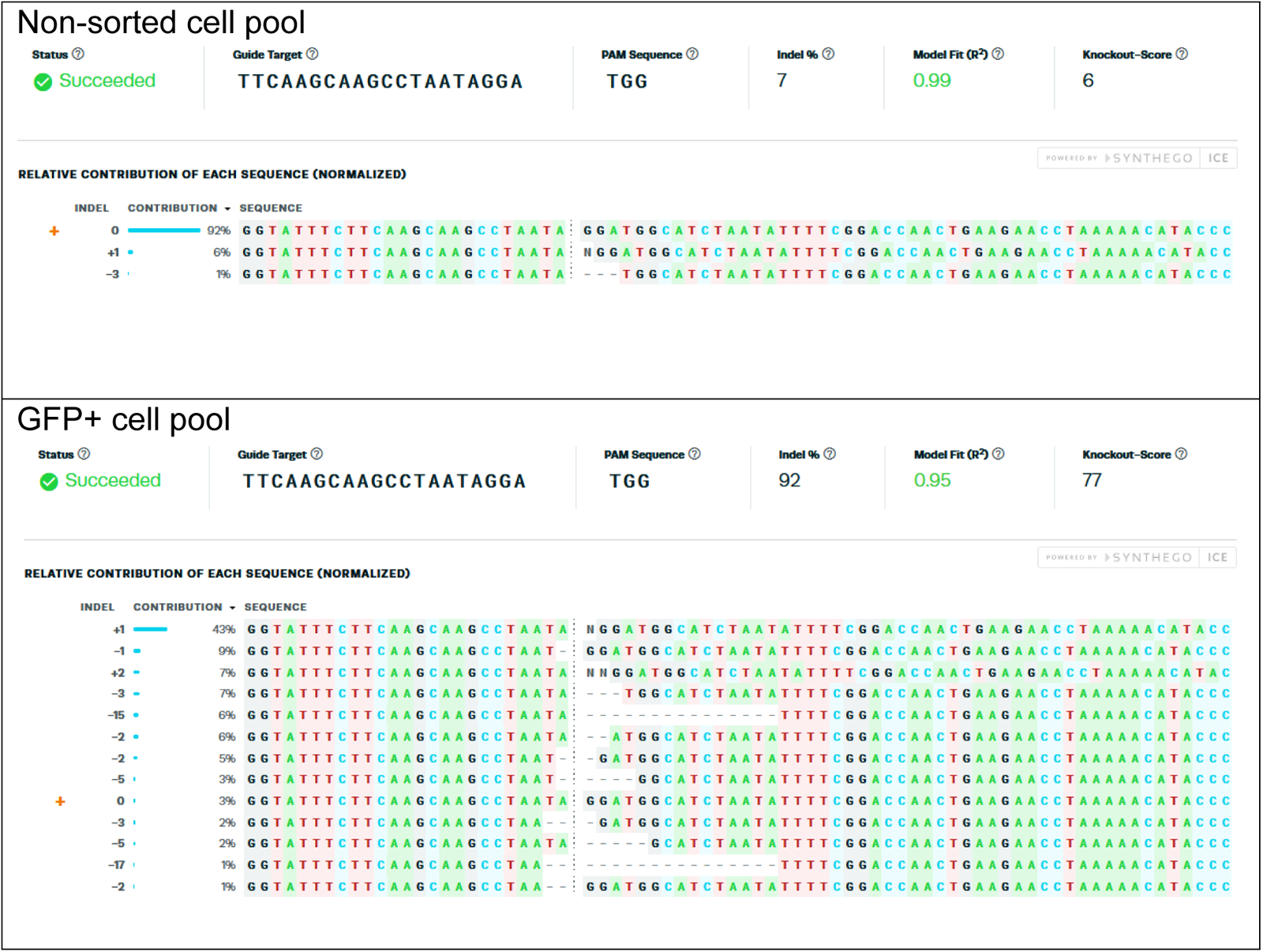
ICE results after amplicon-based cell pool analysis. ICE results provide an overview of the INDEL score, Quality Score (R^2^) and knockout score. In addition, the INDEL distribution is illustrated, showing the proposed sequence. Non-sorted cell pool reveals 92% of the WT sequence, whereas the GFP+ cell pool only shows 3% WT sequence.

**S3 Fig.**
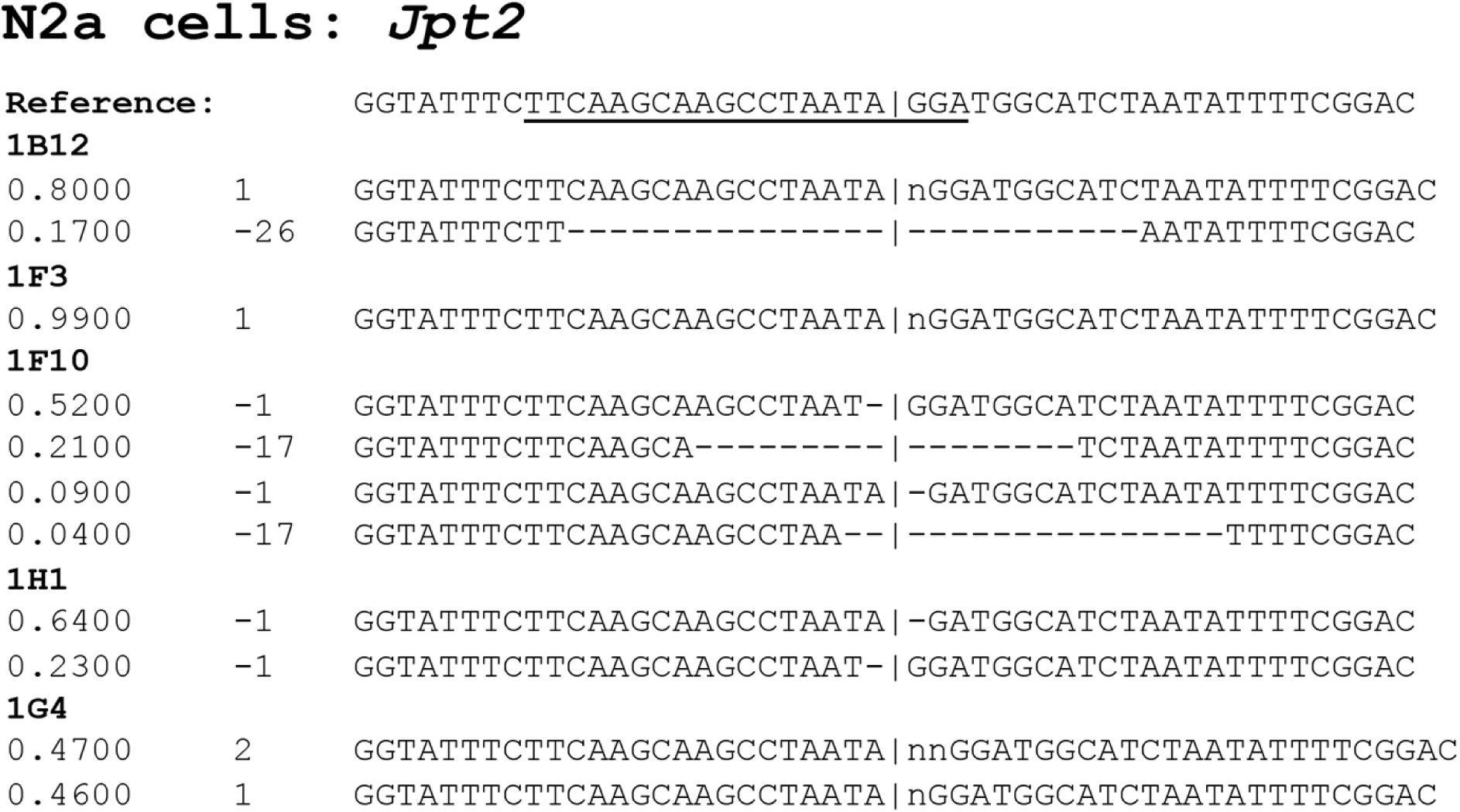
Indels proposed by ICE analysis of *Jpt2* targeted N2a clones. Five high-potential clones and their respective Indels as detected by ICE analysis are shown.

**S4 Fig.**
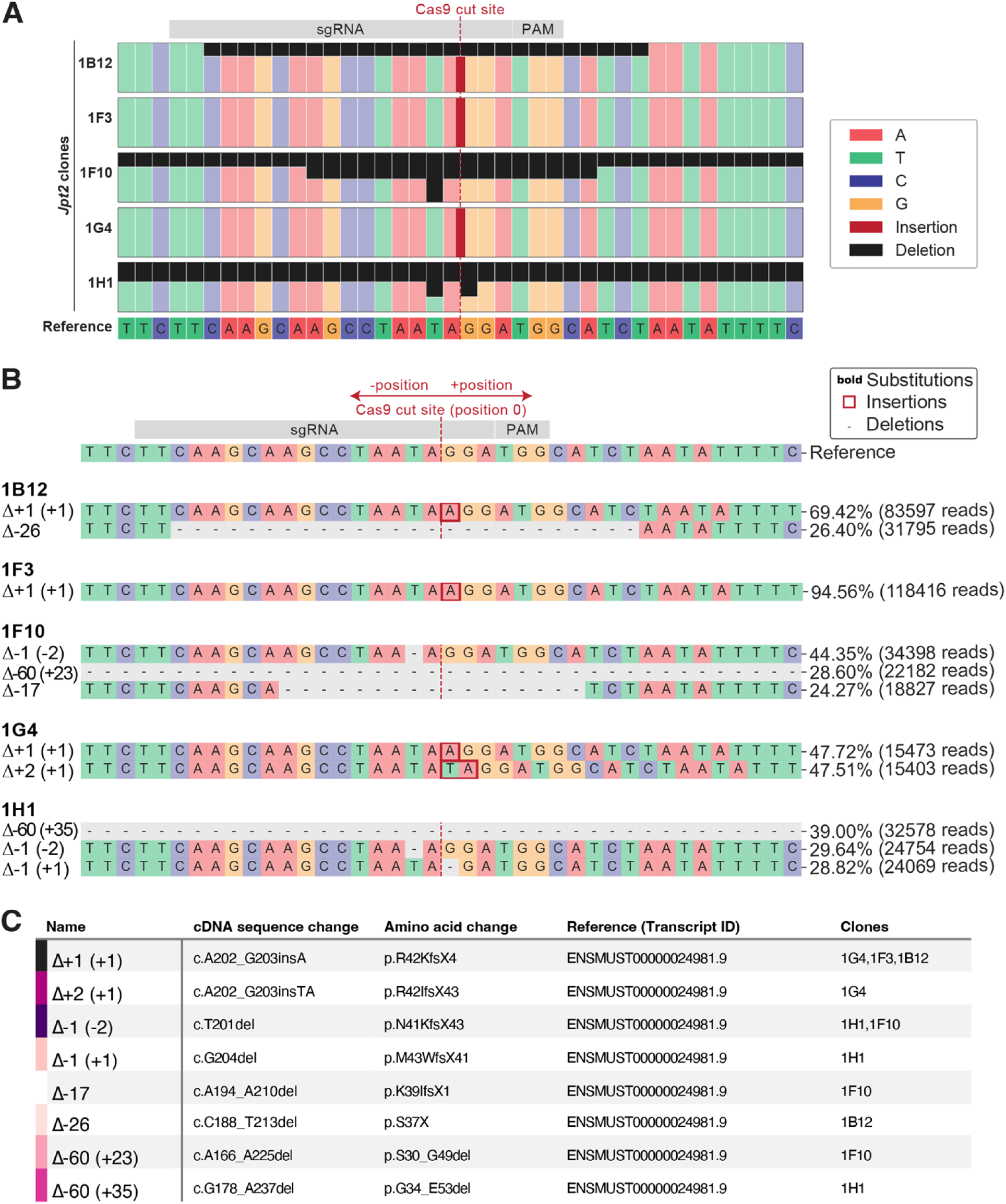
NGS analysis of *Jpt2* N2a cell clones. (**A**) Comparison of identified sequences with highlighting of the position of deleted nucleotides (black) and position of insertions (red). (**B**) Detailed description of the different indels detected by NGS sequencing in the respective clones. (**C**) Overview of the different detected Indels in all *Jpt2* clones, including nomenclature and the clones they are present in.

**S5 Fig.**
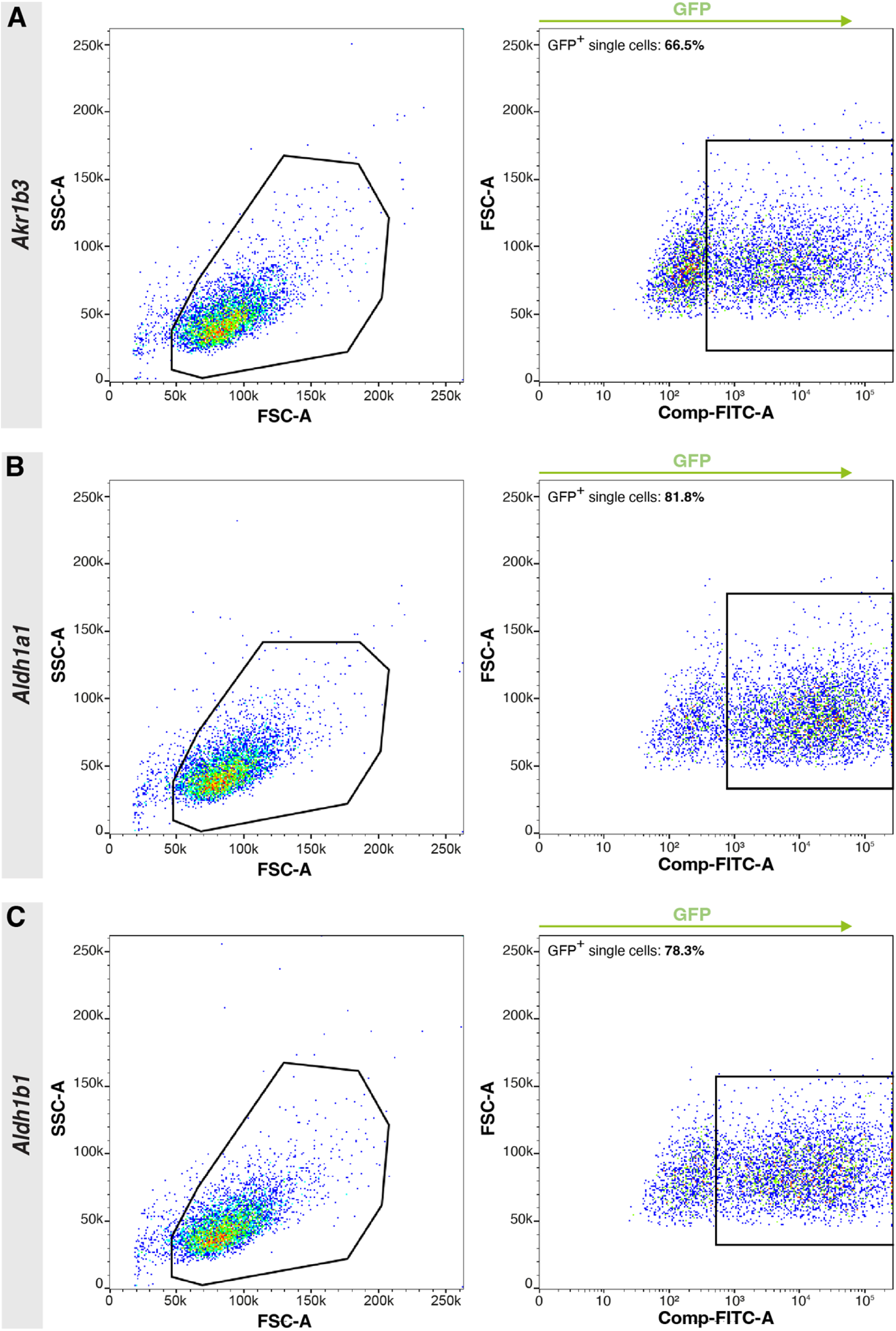
GFP-based FACS sorting of CRISPR/Cas9 treated MCECs. (A) FACS sorting of *Akr1b3* (A), *Aldh1a1* (B) and *Aldh1b1* (C).

**S6 Fig.**
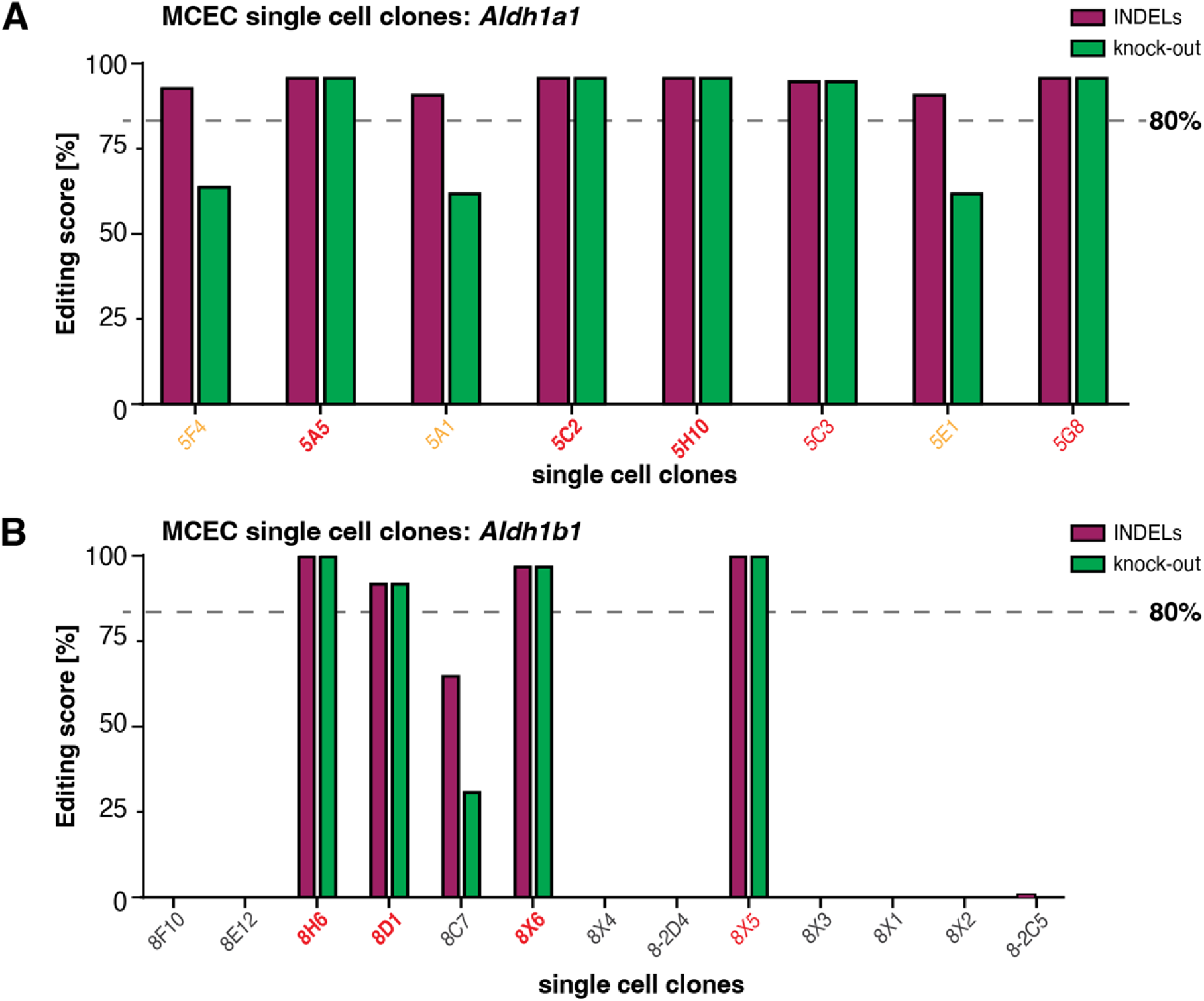
ICE-based screening of *Aldh1a1* and *Aldh1b1* single cell-derived MCEC colonies. (**A**) ICE screening of 8 single cell clones after targeting the *Aldh1a1* locus in MCECs. All colonies met the required quality score of 80%. Clones 5A5, 5C2 and 5H10 were further selected for deep-sequencing analysis (**Fig 3C**). (**B**) ICE screening of *Aldh1b1* MCEC single-cell colonies. 9 out of 13 screened clones showed no sign of editing while 4 out of 14 showed high potential. Clones 8H6, 8D1 and 8X6 were selected for deep-sequencing (**Fig 3C**).

**S7 Fig.**
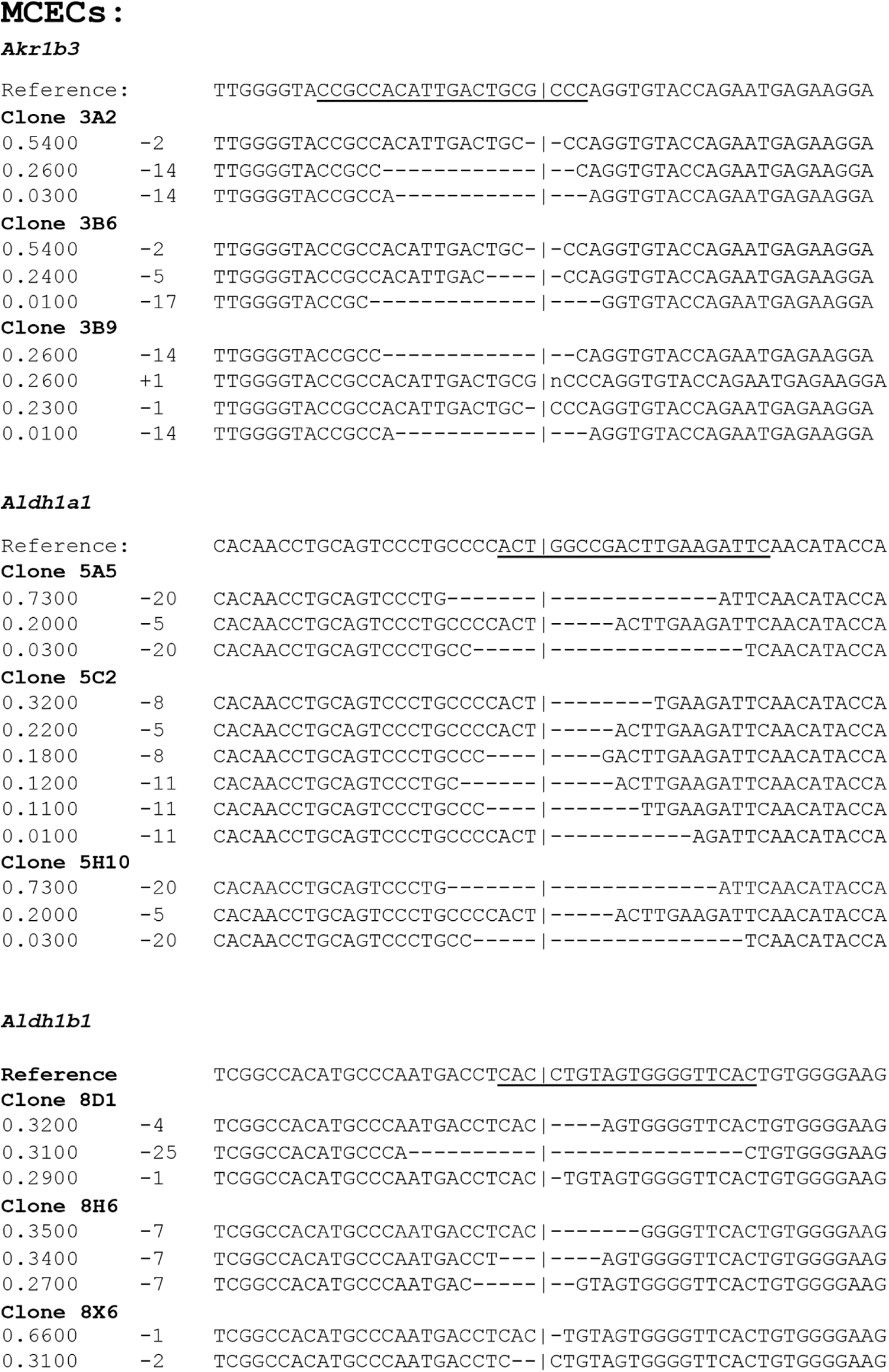
Indels proposed by ICE analysis of MCEC clones. Displays the different target regions for the generated *Akr1b3*, *Aldh1a1* and *Aldh1b1* as analysed using the ICE software. Three items of information are depicted: the Indel distribution (left), the indel size (middle) and the postulated sequence itself (right). *Note:* For *Aldh1a1* and *Aldh1b1,* the reverse complement sgRNA target sequence is displayed (compare to S9 and S10 Figs.).

**S8 Fig.**
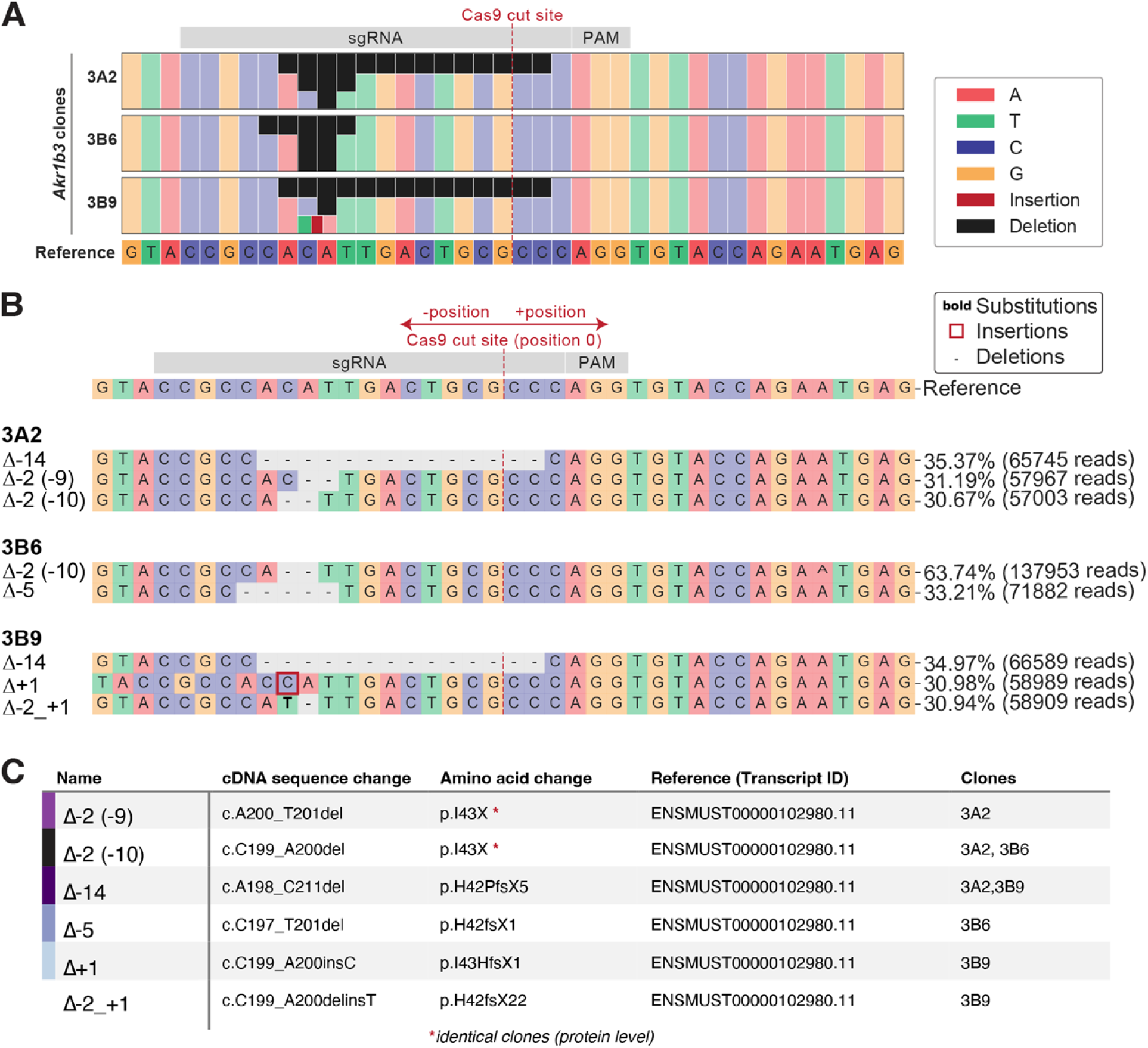
NGS analysis of *Akr1b3* MCEC clones. (**A**) Comparison of identified sequences with highlighting of the position of deleted nucleotides (black) and position of insertions (red). (**B**) Detailed description of the different indels detected by NGS sequencing in the respective clones. (**C**) Overview of the different detected Indels in all *Akr1b3* clones, including nomenclature and the clones in which they are present.

**S9 Fig.**
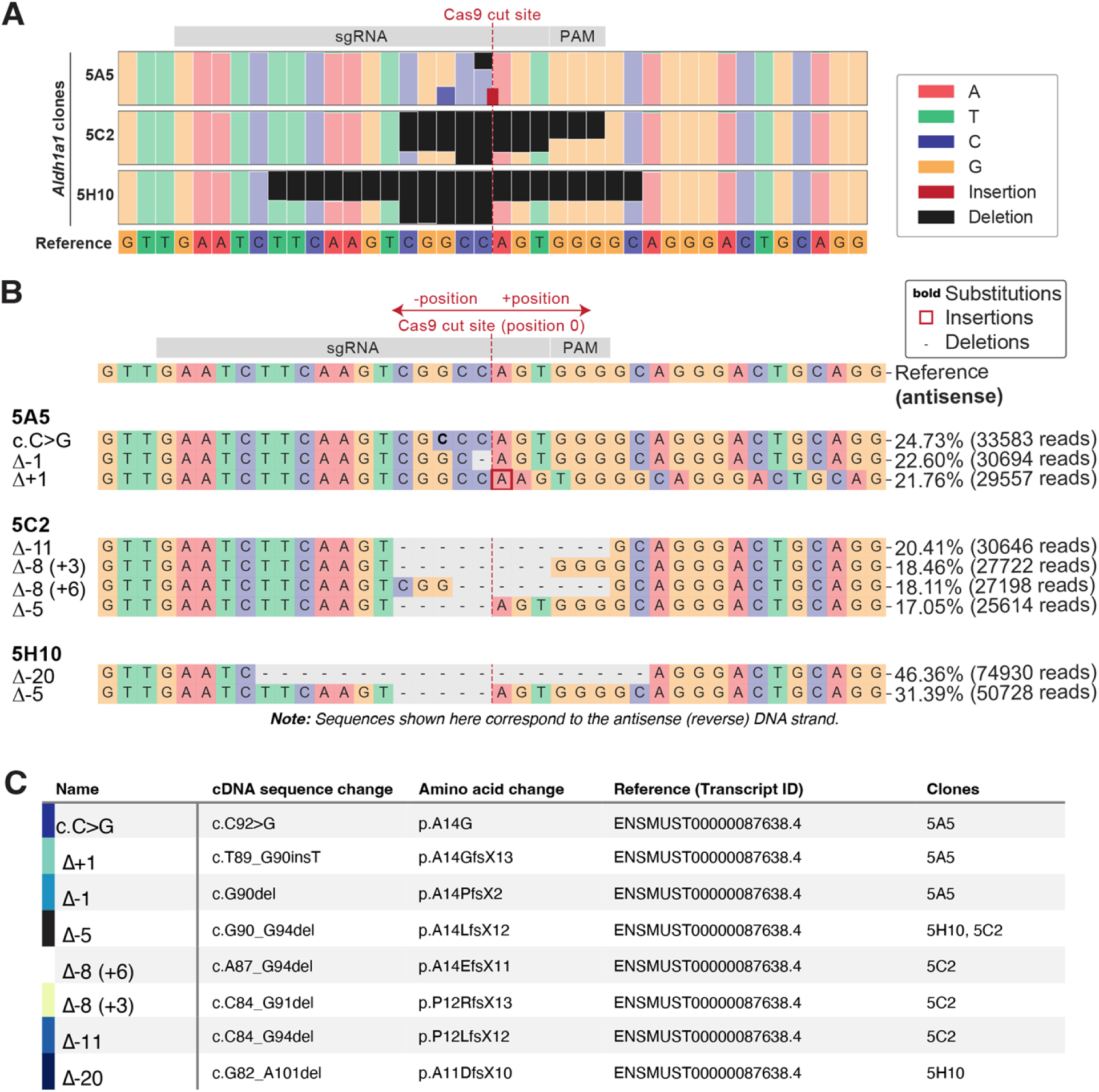
NGS analysis of *Aldh1a1* MCEC clones. (**A**) Comparison of identified sequences with highlighting of the position of deleted nucleotides (black) and position of insertions (red). (**B**) Detailed description of the different indels detected by NGS sequencing in the respective clones. (**C**) Overview of the different detected Indels in all *Aldh1a1* clones, including nomenclature and the clones in which they are present.

**S10 Fig.**
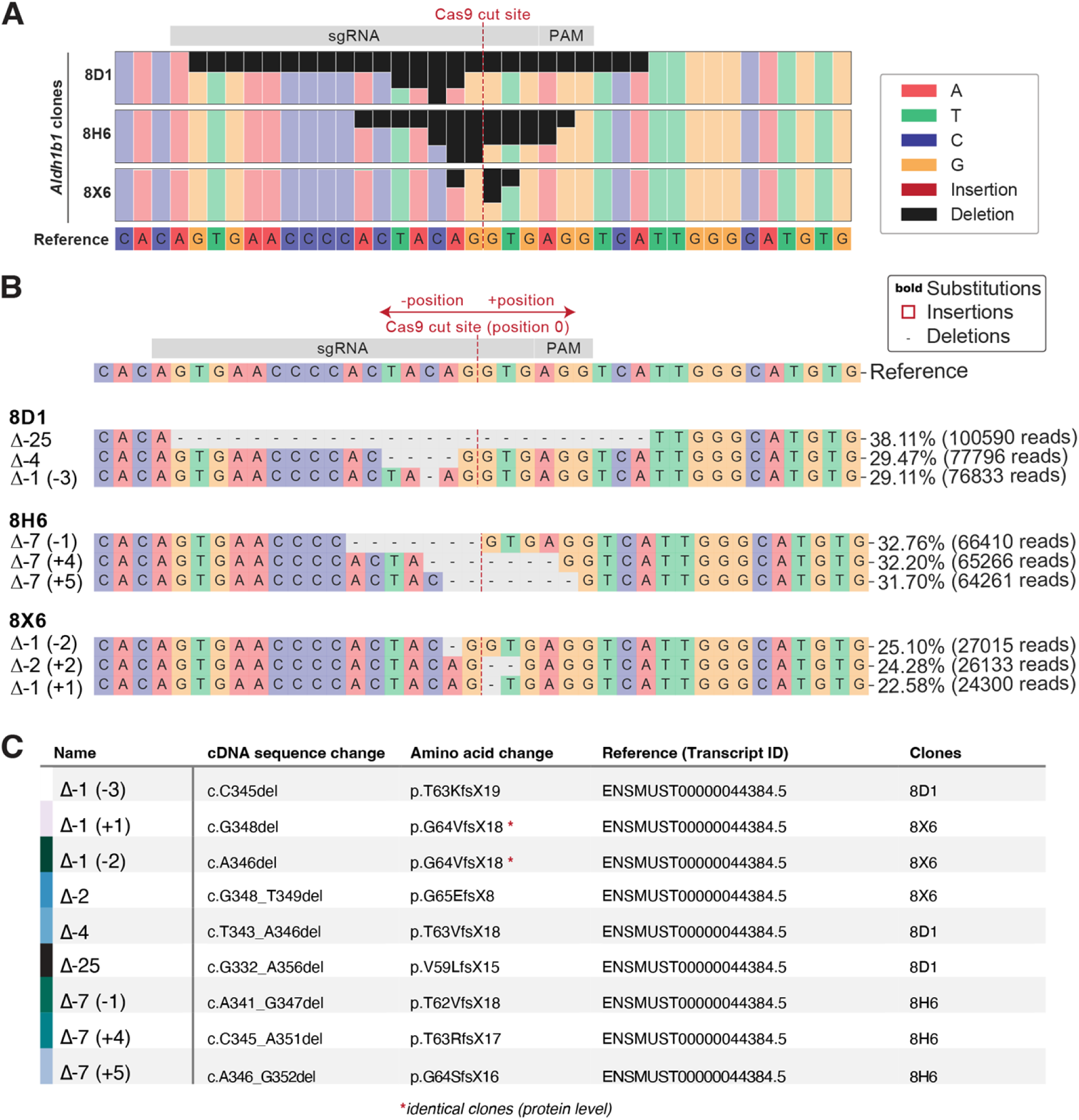
NGS analysis of *Aldh1b1* MCEC clones. (**A**) Comparison of identified sequences with highlighting of the position of deleted nucleotides (black) and position of insertions (red). (**B**) Detailed description of the different indels detected by NGS sequencing in the respective clones. (**C**) Overview of the different detected Indels in all *Aldh1b1* clones, including nomenclature and the clones in which they are present.

